# Pedigree-reconciled pan-genomics reveals megabase-scale hitchhiking after a century of canola breeding

**DOI:** 10.64898/2026.07.29.740455

**Authors:** NC White, KK Gagalova, TE Newman, Y Khentry, LG Kamphuis, MC Derbyshire

## Abstract

Canola breeding has been shaped by strong selection for oil quality, yet the origins of the known oil quality alleles and genomic consequences of their selection are not fully resolved. By integrating pedigree reconstruction with graph pan-genomics we trace inheritance of ancestral genomic regions across historical and contemporary germplasm. Surrounding the low erucic acid allele in *BnA08.FAE1*, we identify a 17.23 Mb haplotype that approached fixation in Australian canola in the early 2000s. Contradicting the prevailing model, this haplotype predates modern breeding and was likely widespread in ancestral *B. napus* in the early 1900s. Genomic analyses implicate centromeric recombination suppression and structural variation in its long-term persistence, which has led to megabase-scale diversity loss through hitchhiking of neighbouring alleles. The haplotype contains extensive structural variation and multiple alleles associated with polygenic disease resistance. Together, these findings reveal the long-term consequences of repeated selection on standing variation during crop improvement.

## Main

Selective breeding over the last century has underpinned the substantial increases in crop production needed to support the world’s growing population^1^. However, crop breeding programs typically focus on a handful of high value traits, such as total yield, seed quality and disease resistance. Intense selection for these traits can severely impact genetic diversity, limiting potential for future crop improvement^2,3^. The consequences of selection may differ markedly between traits controlled by a few major genetic loci and complex polygenic traits.

Hitchhiking around strongly selected loci can substantially reduce local genetic diversity, particularly where recombination is limited^4–6^. For instance, regions surrounding centromeres often undergo limited recombination due to epigenetic suppression and genomic structural variation^7,8^. Heavy selection of loci close to these regions may increase local diversity loss. Diversity loss in crop breeding is also caused by extreme bottlenecks from the use of a limited pool of pedigree founders ^2,9,10^. Haplotype sharing between founders can exacerbate local diversity loss by lowering effective recombination^11^.

Despite these challenges, diversity loss and its causes in selective breeding are under-researched. *Brassica napus* is an ideal species for exploring the genomic impacts of intensive selective breeding. It is the second-most widely grown oilseed and is supported by extensive species-wide genomic data^12^. Furthermore, the pedigrees of many modern *B. napus* cultivars are openly available^13,14^, providing an opportunity to understand the origins and trajectories of alleles important to modern breeding.

The *B. napus* type ‘canola’ is defined by its low seed glucosinolate and erucic acid (EA) content, traits established through intense selection in the 1950s-60s^15,16^. The low EA (LEA) phenotype is traditionally attributed to a single German variety, ‘Liho’, which was used in early Canadian breeding programmes to generate the foundational crosses leading to modern canola^17–21^. Early studies document EA content variability within the original Liho material, suggesting seedstock genetic heterogeneity. Self-propagated zero EA Liho selections were subsequently crossed with locally adapted “Argentine” landraces to establish the LEA trait^20^.

The LEA trait is primarily controlled by two loci containing the homoeologous fatty acid elongase genes *BnaA08.FAE1* and *BnaC03.FAE1*^21–23^. Loss-of-function variants in these genes disrupt erucic acid biosynthesis leading to the canola oil quality phenotype. The S282F substitution in *BnaA08.FAE1* consistently shows the strongest association with EA content in population-wide genomic analyses^24–28^.

Intense and sustained selection for low erucic acid has led to local diversity loss surrounding *BnaA08.FAE1* on chromosome A08^26^. This provides a clear example of diversity loss driven by selection on a major-effect locus. The consequence of this is evident in the linkage drag impacting nearby agronomic traits^29,30^. However, the extent of this diversity loss, its underlying causes, and the origin of the selected LEA haplotypes remain poorly understood.

Contrasting the LEA trait, resistance to *Sclerotinia* stem rot is highly polygenic, challenging conventional breeding^31–34^. This disease, caused by the fungus *Sclerotinia sclerotiorum*, significantly impacts global canola yield^34,35^. Polygenic traits are particularly vulnerable to diversity loss through the gradual erosion of favourable minor-effect alleles through drift, bottlenecks and hitchhiking around loci intensely selected for other traits.

Recent advances in genomics have created new opportunities to characterise genetic diversity in crops. Graph pan-genomics enables representation of sequence and structural variation across multiple high-quality genomes, improving variant genotyping and analysis of complex polymorphisms across large populations^36,37^.

Australian canola breeding is uniquely favourable to investigation of these processes. It has a well-documented pedigree spanning ∼35 years of modern breeding^14^. Through its major founders, this pedigree links isolated Japanese and Canadian breeding programmes, which trace back to early landraces. In this study, we join these three pedigrees and integrate them with graph pan-genomic and phenotypic data to study the impacts of modern breeding on genetic diversity in *B. napus*.

We show that the LEA allele in *BnaA08.FAE1* is embedded in a 17.23 Mb haplotype that originates prior to modern breeding. This haplotype is highly enriched in canola and approached fixation in Australian cultivars in the early 2000s. Contrary to the prevailing model of the LEA trait origin, this haplotype was drawn from widespread standing variation that was repeatedly captured and selected over the last century. Genomic analyses indicate that its persistence is likely driven by recombination suppression in a structurally complex region spanning the A08 centromere.

In parallel, we observe genome-wide erosion of polygenic *Sclerotinia* stem rot resistance in Australian canola, consistent with loss of adaptive variation present in Japanese founders. We identify *Sclerotinia* stem rot resistance QTLs within the LEA haplotype and show that alternative haplotypes at this region were largely lost during Australian breeding. This suggests that oil-quality selection may have restricted access to variation impacting polygenic disease resistance. Together, these findings reveal contrasting but subtly linked modes of diversity loss in breeding. More broadly, this study provides a framework for dissecting the origins and consequences of diversity loss in modern crop improvement.

### Reconciling relationships across a century of global *Brassica napus* breeding

To establish the global ancestry of modern *B. napus*, we first reconciled fragmented historical pedigree records from three national breeding efforts. This uncovered major global bottlenecks in Canadian and Japanese breeding programmes, propagated to Australian canola.

Canadian breeding (1940s-1970s) relied heavily on the Argentine landrace Golden, which was combined with European landraces Liho and Bronowski to produce LEA and double-low cultivars. Accordingly, among 20 Canadian cultivars, 85% were descended from Golden, 75% Liho and 70% Bronowski (Figure 1, Supplementary Table 1). Argentine landrace Nugget is ancestral to just 25%, including the first LEA cultivar Zephyr.

**Figure 1.**
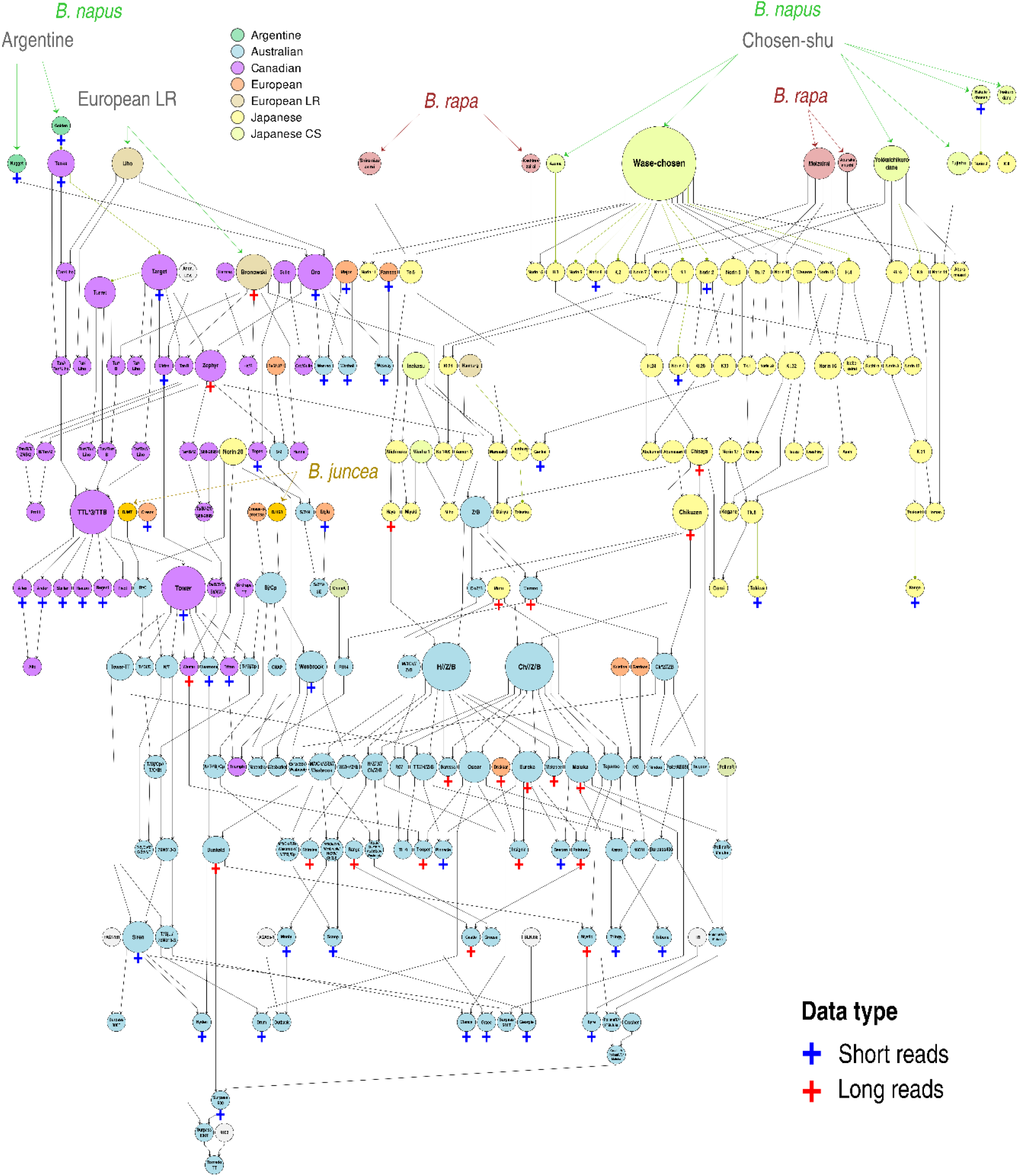
Pedigree combining Australian, Japanese and Canadian pedigrees, coloured by variety type; crosses indicate sequencing technology.

Japanese breeding (1930s-1970) combined *B. napus* with introduced and native *B. rapa*, grown in Japan for ∼400 years. It relied heavily on the European landrace Chosenshu introduced in the late 1800s^13^. This landrace was described at the time as having high resistance to *Sclerotinia* stem rot^13^. Among 60 Japanese cultivars, Chosenshu selections Wasechosen and Yokkaichikurodane are ancestral to 78% and 43%, respectively (Figure 1, Supplementary Table 1).

Beginning in 1970, Australian breeding relied heavily on cultivars from Japanese and Canadian pedigrees. Through extensive use of a few related founders, 86-98% of Australian cultivars descend from Liho, Bronowski, Golden and Nugget, and >70% from Yokkaichikurodane and Wasechosen (Figure 1).

We combined the pedigree with a pan-genome graph comprising 31 genomes (22 new long-read assemblies and 9 references) and short read genotypes called against it for 398 global accessions (Supplementary Table 2 and 3). We called 2,137,846 quality-filtered variants against the complete Xiang5A (X5A) reference genome ^38^; after filtering, 785,160 (including 656,908 SNPs) were retained across 412 samples. Genotyped varieties included 65 from the pedigree, with several cultivars derived from complete sets of sequenced founders (Figure 1).

SNP-and pedigree-based kinship estimates were correlated (*r* = 0.48, P = 0), with an average per-line correlation of 0.48 and ≥0.35 in 74% of lines (Figure 2 A and B), suggesting congruence between genomic data and pedigree records. Principal component analysis (PCA) supported the pedigree structure, separating Japanese and Canadian cultivars along PCs 1-3 (12.3%, 7.9%, and 4.8% variance explained; Figure 2 C and Supplementary Figure 1). Argentine landraces Nugget and Golden clustered with Canadian lines along PC1, with PCs 2 and 3 resolving sub-structure around these founders (Figure 2 C and Supplementary Figure 1).

**Figure 2.**
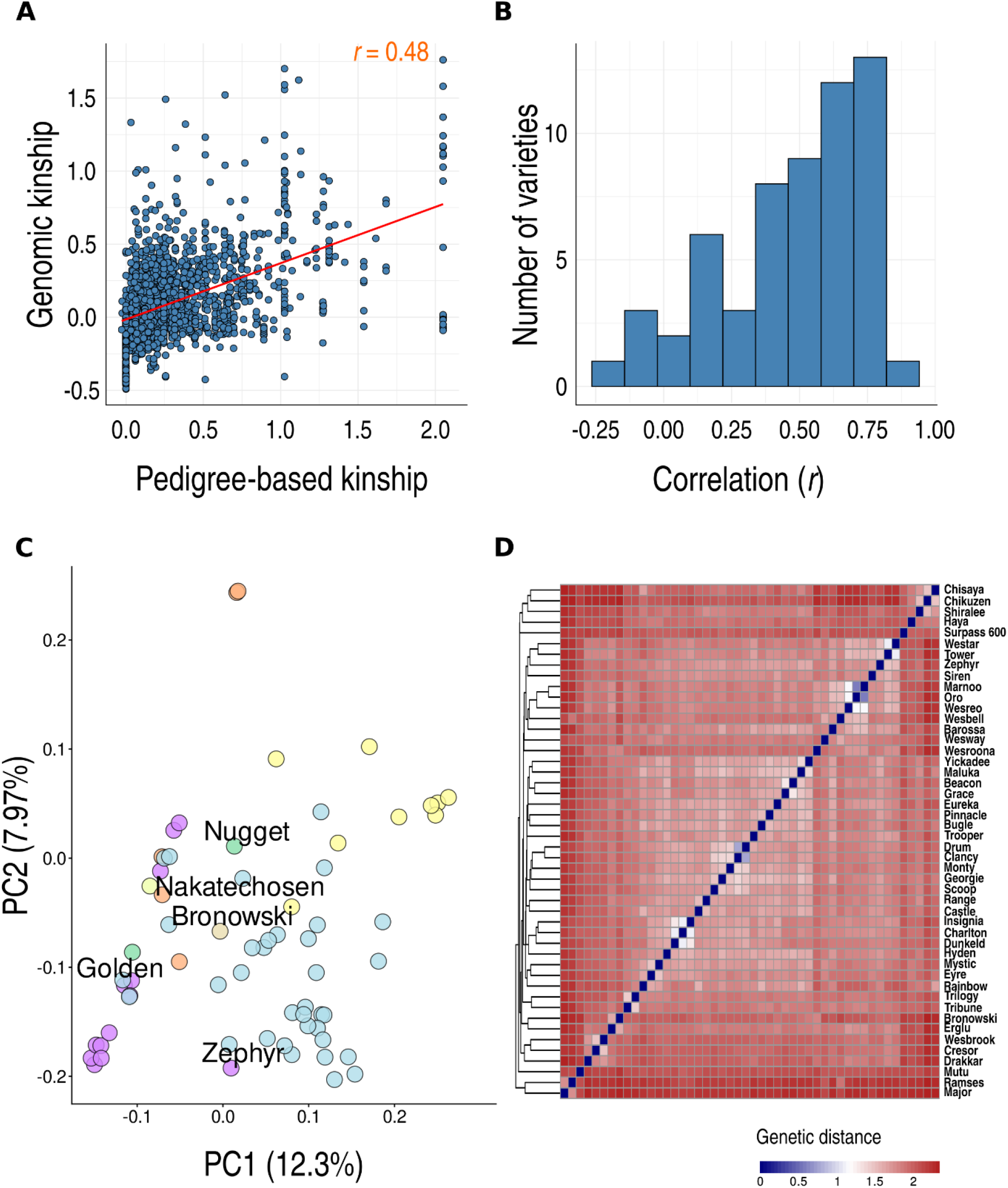
**A** Correlation between genomic (y) and pedigree (x) based kinship. **B** Histogram of correlations between genomic and pedigree-based kinship for each cultivar paired with all others (bins = 10). **C** principal component analysis of genome-based kinship; points coloured by variety type as for Figure 1. **D** Heatmap showing distances derived from genome-based kinship between Australian cultivars estimated with base allele frequencies derived from known founders shows the inverse of the coefficient of kinship^39^.

Consistent with their mixed ancestry, Australian cultivars captured the greatest diversity (9.0% of variance), three times that of Canadian and Japanese cultivars (2.7%). They spanned the genetic space between these groups along PCs 1 and 2 (Figure 2 C), with additional separation across PC3 alongside European material, likely reflecting less extensive European introgression (Supplementary Figure 1). Clear congruence between genomic data and pedigree was found in the close relationships between some Australian cultivars, such as the siblings Drum and Clancy, and Charlton and its descendants Hyden, Mystic and Eyre (Figure 2 D).

Overall, this pedigree-resolved graph pan-genome spanning a century of *B. napus* breeding provides a resource for high value trait introgression tracking and can be overlayed with additional genome sequencing and pedigree data in future.

### Intense oil quality selection trapped a 17 Mb structural haplotype on chromosome A08

Having established the global ancestry of modern canola, we next investigated genomic consequences of selection. Selection scans across Australian cultivars identified a 17.23 Mb unbroken haplotype encompassing *BnaA08.FAE1* (9.28 to 26.5 Mb in X5A; Figure 3 A). This haplotype had close to zero diversity in Australian cultivars (n = 34), contrasting the higher diversity in Australian pedigree founders and surrounding genomic regions (Figure 3 B). The haplotype is in approximately half of Canadian cultivars and almost fixed in Australian cultivars (30/34 lines; Figure 3 C).

**Figure 3.**
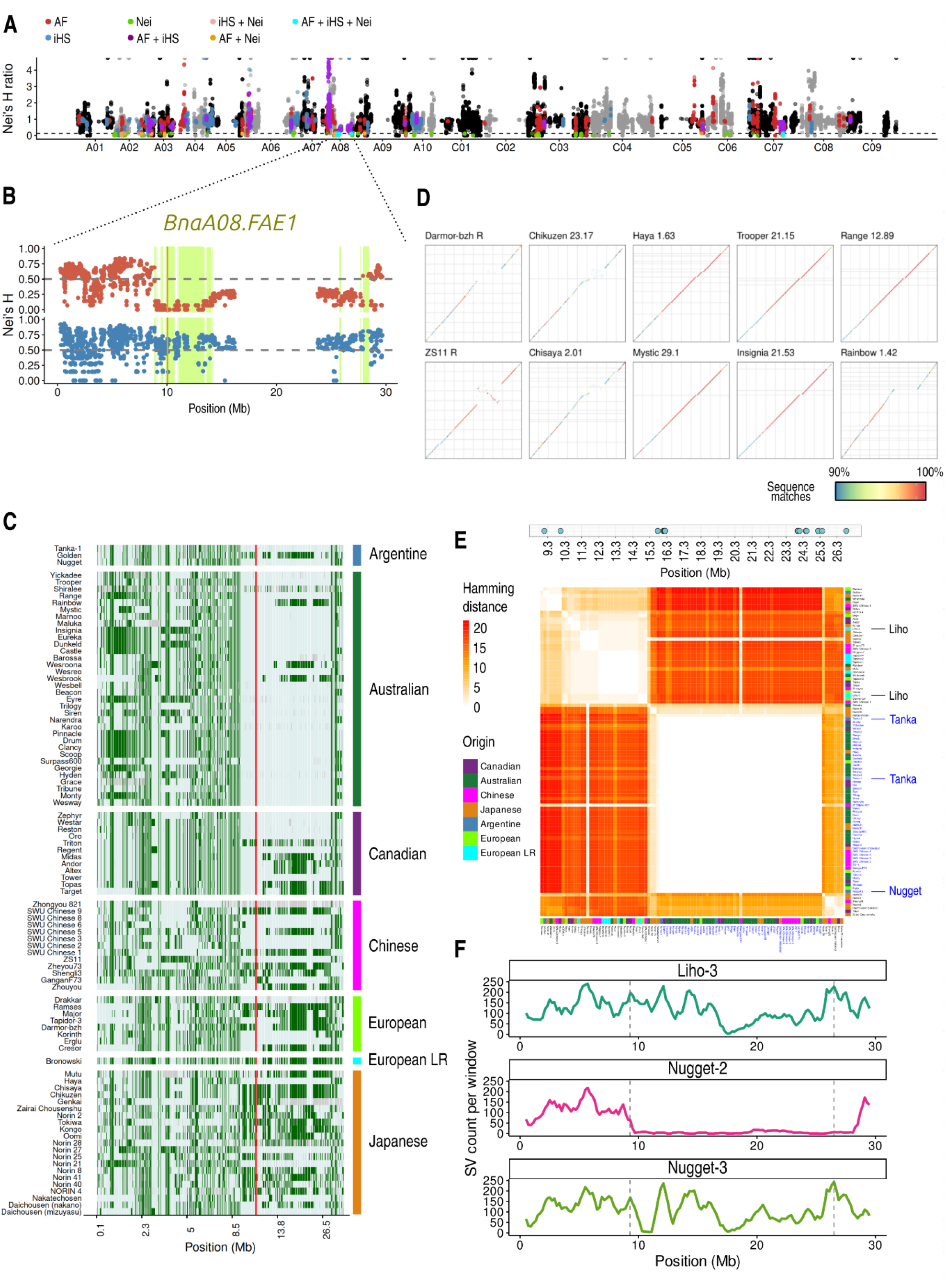
Selection for oil quality across a 17 Mb haplotype containing a low erucic acid allele on A08. **A** Results of selection scans in Australian cultivars based on aberrant allele frequency (AF), integrated haplotype score (iHS), and Nei’s haplotype diversity difference between Australian pedigree founders and descendant cultivars (Nei). Colours represent regions meeting a 95^th^ percentile threshold in one or more test. **B** Nei’s haplotype diversity in 100 Kb sliding windows across genomic coordinates (x axis) in the X5A genome. Blue points at the bottom are for 14 major Australian pedigree founders and red points at the top are for 34 cultivars from the same pedigree. Green shaded areas met the 95^th^ percentile threshold for Nei’s haplotype diversity difference. The vertical gold line is the *BnaA08.FAE1* locus. **C** Haplotypes across A08-17-LEA for selected lines sequenced in this study. Lines include those from the global pedigree and additional Japanese and Chinese lines for further context. Light blue is reference and green is the alternate allele. **D** Synteny with X5A across A08. Assembly N50s are above plots and “R” denotes a downloaded chromosome-scale reference assembly. Colouring shows percent of matching bases for each alignment. **E** Hamming distance across 23 SNPs callable with two Liho accessions from NCBI SRA. Positions of SNPs across A08 are given above in the plot with blue points. **F** Structural variants (y axis) called against X5A across 100 Kb sliding windows for Liho and Nugget. Further information on SRA accessions is in Supplementary Table 4.

Long-read assemblies confirmed that this haplotype contains the LEA S282F allele in *BnaA08.FAE1* (Supplementary Figure 2). Unexpectedly, the haplotype was found in Argentine-selections Nugget and Tanka (Figure 3 C). Its absence in Golden, Tanka’s parental selection, suggests heterogeneity within Argentine landraces that founded the Canadian pedigree. Its presence in these landraces suggests it was in *B. napus* standing variation before modern breeding. Supporting this, it was also found in four pre-1970 Japanese cultivars not selected for LEA (Figure 3 C), and six Chinese cultivars. We refer to this region as “A08-17-LEA”.

This haplotype is in the chromosome-scale reference genome for X5A. Alignment of long reads contigs indicated 100% identity and synteny among carrier genomes, supporting identity by descent (IBD) (Figure 3 D). To investigate the origin of this haplotype, we analysed multiple accessions labelled as “Liho”, the presumed donor of the *BnaA08.FAE1* S282F allele. Unexpectedly, A08-17-LEA was not found in any of them. Across 23 SNPs distributed throughout A08-17-LEA, two Liho short read accessions clustered away from A08-17-LEA carriers, including an additional Tanka short read accession (Figure 3 E).

Long read data from a third Liho accession confirmed this as it showed extensive structural variation relative to X5A. Surprisingly, the *BnaA08.FAE1* S282F substitution was not in the Liho long read data (Supplementary Figure 2). However, though it wasn’t called due to low coverage in one short read accession, it was detected in the other (0/0 call at A08 position 10048887; Supplementary Table 5).

Further supporting an occurrence in founding Argentine landraces, long reads from an accession labelled as “Nugget” showed only 2 structural variants across the entire 17 Mb of A08-17-LEA in contrast to the 447 of Liho (Figure 3 F). A second Nugget long reads accession contained 263 structural variants, further supporting heterogeneity in founding Argentine strains. Notably, both Nugget long read accessions contained the *BnaA08.FAE1* S282F allele (Supplementary Figure 1), which coincided with a loss of structural variation either side of the gene (Figure 3 F).

We propose an alternative model for the origin of the LEA allele on A08. Our data suggest that this allele is embedded in an extended haplotype that was present in *B. napus* standing variation before modern breeding. This may have been indirectly selected from segregating Argentine seedstocks after crosses with Liho in Canada and in some programmes from varieties related to east Asian germplasm. The near fixation of A08-17- LEA in Australian canola reflects repeated capture of a common pre-existing haplotype rather than the spread of a rare or novel mutation as previously suggested. These findings show how selection for a single high value trait can drive near fixation of a large haplotype, almost eliminating diversity across more than half of a chromosome.

### Centromeric recombination suppression and selection of common standing variation may underpin severe diversity loss on A08

We next investigated why A08-17-LEA has remained intact despite multiple generations of recombination. We identified 165 unique haplotypes across the region containing A08- 17-LEA among 412 diverse global *B. napus* accessions (Figure 4 A). We found that A08- 17-LEA (>99.999% SNP identity with X5A) was widespread, occurring in 44% of accessions. This supports an origin predating modern breeding rather than recent pedigree connectivity.

**Figure 4.**
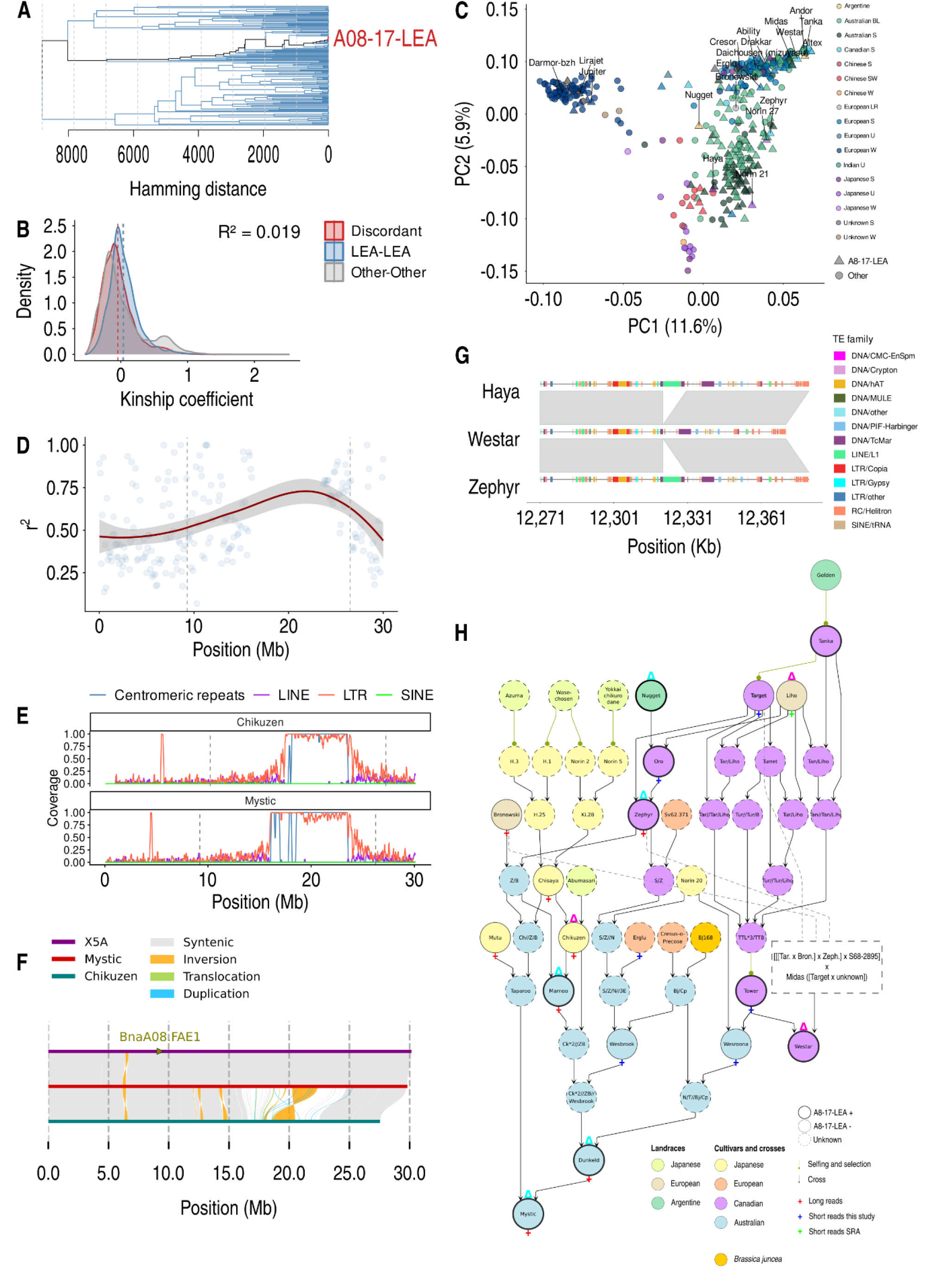
**A** Dendrogram showing relationships between 165 unique haplotypes across A08-17-LEA. **B** The x axis shows kinship coefficients between pairs where both are A08- 17-LEA carriers (LEA-LEA), one is and the other is not (Discordant) and both are not carriers (Other-Other). **C** Principal component analysis based on genome-wide kinship, where different colours represent different breeding programmes and types (S = spring; W = winter; SW = semi-winter; LR = landrace; U = unknown; BL = breeding line); shapes represent whether lines are A08-17-LEA carriers or not. **D** Linkage disequilibrium (r^2^) on the y axis across 100 Kb sliding windows for Japanese cultivars across A08. **E** Number and type of repeat elements in 100 Kb sliding windows across A08 for A08-17-LEA carrier Mystic and non-carrier Chikuzen; vertical dashed line represents limits of the A08-17-LEA region on chromosome A08 based on alignment to X5A. **F** Alignment between carriers X5A, Mystic and non-carrier Chikuzen; the position of *BnA08.FAE1* is marked out in gold; ribbons show structural rearrangements. **G** Insertion of a LINE/L1 element in A08-17-LEA carriers Haya and Zephyr relative to carrier Westar. Grey ribbons show synteny and features are coloured based on transposon family (TE family). **H** The pedigrees of Australian cultivar Mystic and Canadian cultivar Westar. Dashed lines and boxes summarise the Westar pedigree, which is omitted to save space. Variety types are illustrated with different colours, alongside shapes and line thickness for sequencing data type, presence of A08-17-LEA and the LINE/L1 insertion from **G**.

To test this, we assessed the association between the distribution of A08-17-LEA and genome-wide relatedness. Presence of A08-17-LEA explained only a small proportion of genome-wide kinship (R^2^ = 0.0187, P < 2e^-16^; Figure 4 B), with PCA showing that A08-17- LEA carriers are dispersed across geographically and genotypically distinct clusters. A logistic regression on the first three PCs explained only modest variance in presence or absence of A08-17-LEA (McFadden’s pseudo-R² = 0.138) (Figure 4 C).

Overall, genome-wide population structure was most strongly associated with geographic origin and growth habit, whereas A08-17-LEA occurred largely independently. For example, Australian pedigree founders Haya (Japanese) and Zephyr (Canadian) are genotypically distant despite both possessing A08-17-LEA. They are distributed either side of most Australian cultivars along both PC1 (11.6% variance) and PC2 (5.9% variance), consistent with their documented use in Australian breeding (Figure 4 C). These observations are inconsistent with seedbank mislabelling or more recent common ancestry of A08-17-LEA carriers.

Persistence of an intact 17 Mb haplotype over numerous breeding generations implies limited recombination. Supporting this, LD peaked within A08-17-LEA across Japanese cultivars with diverse A08 sequences (Figure 4 D). A strong enrichment of centromeric repeats and long terminal repeat (LTR) elements within A08-17-LEA in high quality assemblies suggests it spans the centromere (Figure 4 E). This indicates that centromeric recombination suppression is a likely mechanism for its long-term maintenance. Multiple large inversions between A08-17-LEA and divergent haplotypes, such as that in Japanese cultivar Chikuzen, could further limit recombination by disrupting chromatid homology (Figure 4 F).

Consistent with a deep ancestry of A08-17-LEA, we also identified a long interspersed nuclear element (LINE) L1 insertion in carriers Zephyr and Haya but not Westar (Figure 4 G). Supporting its ancient origin, the insertion was also in Nugget, multiple Australian cultivars derived from Zephyr and Haya and the French carrier Tapidor. Though Westar is partially descended from Nugget, it has more extensive ancestry from Tanka (Figure 4 H), suggesting it may carry a different ancestral copy of A08-17-LEA.

Supporting this, Westar also contained a hAT insertion relative to both Zephyr and Haya. This insertion was in Australian cultivars Trooper, which is descended from Westar, and Eureka and its descendants Dunkeld and Mystic (Supplementary Figure 3). Since Eureka descends only from Zephyr and Japanese cultivars, this could indicate its inheritance of the Tanka haplotype via residual segregation in Zephyr.

Illustrating the structural stability of A08-17-LEA, it has been retained in the Australian cultivar Mystic despite the extensive historical crosses and diverse ancestry involved in its development (Figure 4 H). The substantial structural variation in A08-17-LEA between Mystic and non-carrier Chikuzen suggests potential functional consequences that remain to be characterised.

Collectively, these results support a model in which A08-17-LEA has been maintained in *B. napus* standing variation due to centromeric recombination suppression and structural complexity. This illustrates how intrinsically low recombination within large haplotypes can drive long-range chromosomal diversity loss during breeding.

### Loss of resistance to a neglected disease may have been an unintended consequence of diversity loss during breeding

Given the extent of diversity loss surrounding A08-17-LEA, we next asked whether selection for it may have reduced standing variation impacting other traits. We focused on polygenic *Sclerotinia* stem rot resistance as it has been documented among some varieties in the Australian pedigree and its Japanese founders^34,40,41^. Furthermore, Japanese pedigree records suggest extensive use of the landrace Chosenshu, which was widely favoured for its *Sclerotinia* stem rot resistance^13^. Lesion length screens from three environments showed moderate pairwise correlations ranging from 0.35 to 0.54 (Figure 5 A) indicating environmental variation and genotype-by-environment effects. Accordingly, narrow-sense heritability (*h*^2^) was relatively low within environments, ranging from 0.26 to 0.35 (Figure 5 B).

**Figure 5.**
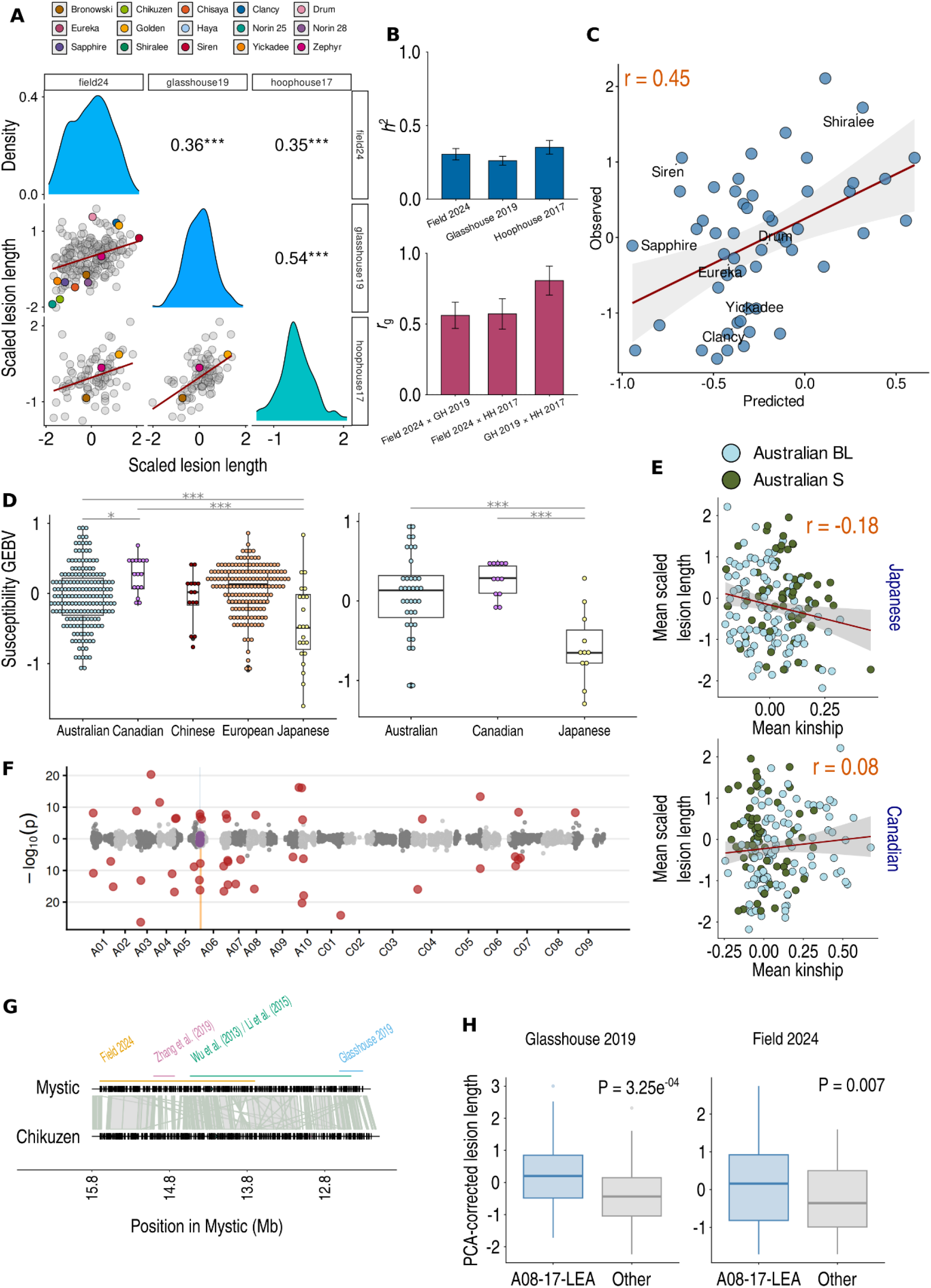
**A** Correlation between lesion lengths in three environments; fta24 = Field 2024, glasshouse19 = Glasshouse 2019^31^, hoophouse17 = Hoophouse 2017. Density plots show the distribution of normalised lesion scores and Pearson’s correlation coefficient to the right; *** = P < 0.0001. **B** Top: narrow sense heritability (h^2^) estimated from a genomic best linear unbiased predictor (G-BLUP) model; bottom: genetic correlation (r_g_) between environments from a G-BLUP model. Error bars show standard error. **C** G-BLUP genomic estimated breeding values (GEBVs) (x) against normalised lesion lengths (y) for Australian cultivars masked from the model; Pearson’s correlation top-left. **D** G-BLUP GEBVs classified based on geographic origin for all varieties (left) and varieties in the global pedigree (right); *** = P < 0.0001, * = P < 0.05; boxes and whiskers show interquartile range. **E** Correlation between normalised lesion length (y) and average kinship (x) to Japanese (top) and Canadian (bottom) for Australian cultivars and breeding lines; Pearson’s correlation coefficient top-right; red line = linear regression of y onto x. **F** Manhattan plot showing QTLs detected in glasshouse (y axis downwards) and field environments (y axis upwards). Red points were detected in one environment and purple points are in linkage disequilibrium blocks containing QTLs from both. **G** A08-17-LEA region surrounding *Sclerotinia* stem rot resistance QTLs from this and two previous studies (colour-coded). Alignment between reverse-complemented extracts with original reversed Mb positions in Mystic on the x axis; ribbons represent alignments and arrows gene annotations. **H** Lesion length corrected for population structure (three principal components) on the y axis, showing a difference between A08-17-LEA carriers and non-carriers (Other). Comparison among Australian and east Asian varieties. Boxes and whiskers show interquartile range and P values are from a linear regression.

Despite moderate phenotypic correlations, pairwise genetic correlations were moderate to high, ranging from 0.56 (+/- 0.09) to 0.81 (+/- 0.10), suggesting relatively stable genomic effects across environments (Figure 5 B). Furthermore, key varieties showed consistent contrasting levels of resistance across environments. For example, Argentine landrace Golden, Canadian cultivar Zephyr and the low glucosinolate founder Bronowski consistently spanned the spectrum of resistance in the same rank order across the three environments (Figure 5 A). Supporting trait environmental stability, genomic best linear unbiased predictor (G-BLUP) models showed a moderate predictive ability (r = 0.45, P < 2.2e^-16^) for Australian cultivars in the field environment when incorporating genotypes and phenotypic data from the other two (Figure 5 C).

High resistance in the Japanese varieties Chisaya and Haya^31,40,42^ and their descendant Australian cultivars Yickadee and Beacon suggests possible inheritance from the Japanese pedigree (Supplementary Figure 5). Supporting this, we found that *Sclerotinia* stem rot resistance breeding values estimated from genomic data were more favourable in Japanese than Canadian varieties, with Australian cultivars intermediate (P = < 2.2e^-16^; Figure 5 D). Furthermore, stem lesion length was negatively correlated with kinship to Japanese (r = −0.18, P = 0.02) but not Canadian (r = 0.08, P = 0.27) cultivars (Figure 5 E).

Confirming the polygenic nature of *Sclerotinia* stem rot resistance, genome-wide association in field and glasshouse^31^ environments detected only minor QTLs (Supplementary Table 6). The most consistent associations were found on chromosomes A01, A06 and A08 (Figure 5 F). Though these associations were only detected in one environment after false discovery rate correction, linear modelling with correction for population structure detected significant effects in both (R^2^ from 1.05-10.53%; Supplementary Table 6).

Underlining the potential impacts of localised diversity loss on phenotypic potential, we found one significant association in each environment within A08-17-LEA (Figure 5 G). The SNP at position 11,556,270 was the strongest association detected, explaining 10.53% variance in the glasshouse (linear model P = 2.32e^-6^; BLINK P adjusted < 0.0001). It showed a consistent directional effect in the field, explaining 1.16% variance (linear model P = 0.034). Further support for the association between A08-17-LEA and *Sclerotinia* stem rot resistance comes from multiple studies reporting QTLs within this region^43–46^ (Figure 5 H). This includes a QTL intersecting the region surrounding position 11,556,270^44,45^.

Among Australian and east Asian varieties, A08-17-LEA carriers were significantly more susceptible after population structure correction than non-carriers in the field (P = 0.007) and glasshouse (P = 3.25e^-4^), supporting an association between this haplotype and resistance (Figure 5 H). Haplotypes lost in Australian breeding, such as the one in partially resistant Japanese cultivar Chikuzen, could be an important but neglected source of diversity for *Sclerotinia* stem rot resistance. Of note was a rapid alkalinisation factor (RALF) gene from Chikuzen missing in A08-17-LEA carrier Mystic within a QTL-dense region (Figure 5 G, Supplementary Table 7). RALFs have diverse roles in plant immunity and have shown a potential role in defence against *S. sclerotiorum*^47^. Further characterisation of this and other defence-associated genes within A08-17-LEA may therefore be of interest.

Overall, our data suggest that substantial heritable *Sclerotinia* stem rot resistance present in Japanese germplasm has been reduced in Australian canola. A lack of selection for this trait, coupled with extensive use of Canadian cultivars with a narrow and relatively susceptible genetic base, may have contributed to this erosion. While effect sizes are small and environment-dependent, the reduction in resistance-associated variation in Australian germplasm is consistent with loss of polygenic resistance during breeding. Localised diversity loss at A08-17-LEA, a region overlapping *Sclerotinia* stem rot resistance QTLs from multiple studies, may have restricted access to haplotypic variation relative to the trait. These associations do not establish causality, but they identify A08-17-LEA as a region where oil-quality selection may have constrained variation relevant to disease resistance. Together, these results link two contrasting modes of diversity loss during breeding: localised hitchhiking around strongly selected loci and gradual erosion of polygenic variation.

## Methods

### Genome sequencing

For Illumina sequencing, seeds were planted in UWA plant biology mix compost (Richgro, Australia) and grown for four weeks. Two hundred milligrams of leaf tissue was sampled, and DNA was extracted using the DNeasy Plant Mini Kit (Qiagen) following manufacturer’s instructions. For long read sequencing, plants were grown in a glasshouse in 24 cell trays using UWA plant biology mix. VectoBac (Valent Biosciences) was applied as a pesticide 9 days after planting. The cultivar “Mutu” failed to germinate during this trial and was thus later grown in a Conviron under long day conditions (16- hour day: 8-hour night) at 21 °C. Three days after thinning, 1.25 g of Nitrophoska Perfect (EuroChem) fertiliser was added. Once four leaves had developed, after four to five weeks, two to three leaves were cut off (equalling about 3 g of total plant matter) and snap frozen in liquid nitrogen. High molecular weight (HMW) DNA was extracted using the Nucleobond HMW DNA kit with a slightly modified protocol for resuspension and homogenisation. The kit was used as per manufacturer’s instructions until final homogenisation steps where the sample pellet was kept overnight at 4 °C in the 50 mL Falcon tube before being heated to 37 °Cfor 60 minutes. The sample was then transferred with a wide-bore pipette tip into a 1.5 ml DNA LoBind Eppendorf tube which was incubated in a thermomixer for two hours at 37 °C and 300 rpm.

### Pedigree construction

Published pedigrees were obtained for Japanese^13^, Canadian^48–50^ and Australian^14^ *B. napus*, spanning a period from approximately 1930 to 2005. These pedigrees were manually reconciled by constructing a simplified input (Supplementary Table 8) for the Helium pedigree plotter^51^. The global pedigree was plot using Helium and adjusted aesthetically for presentation.

### *Sclerotinia* stem rot resistance screens

*Sclerotinia* stem rot resistance data from three environments were used. Replicates are individual inoculated plants (Table 1).

**Table 1.**
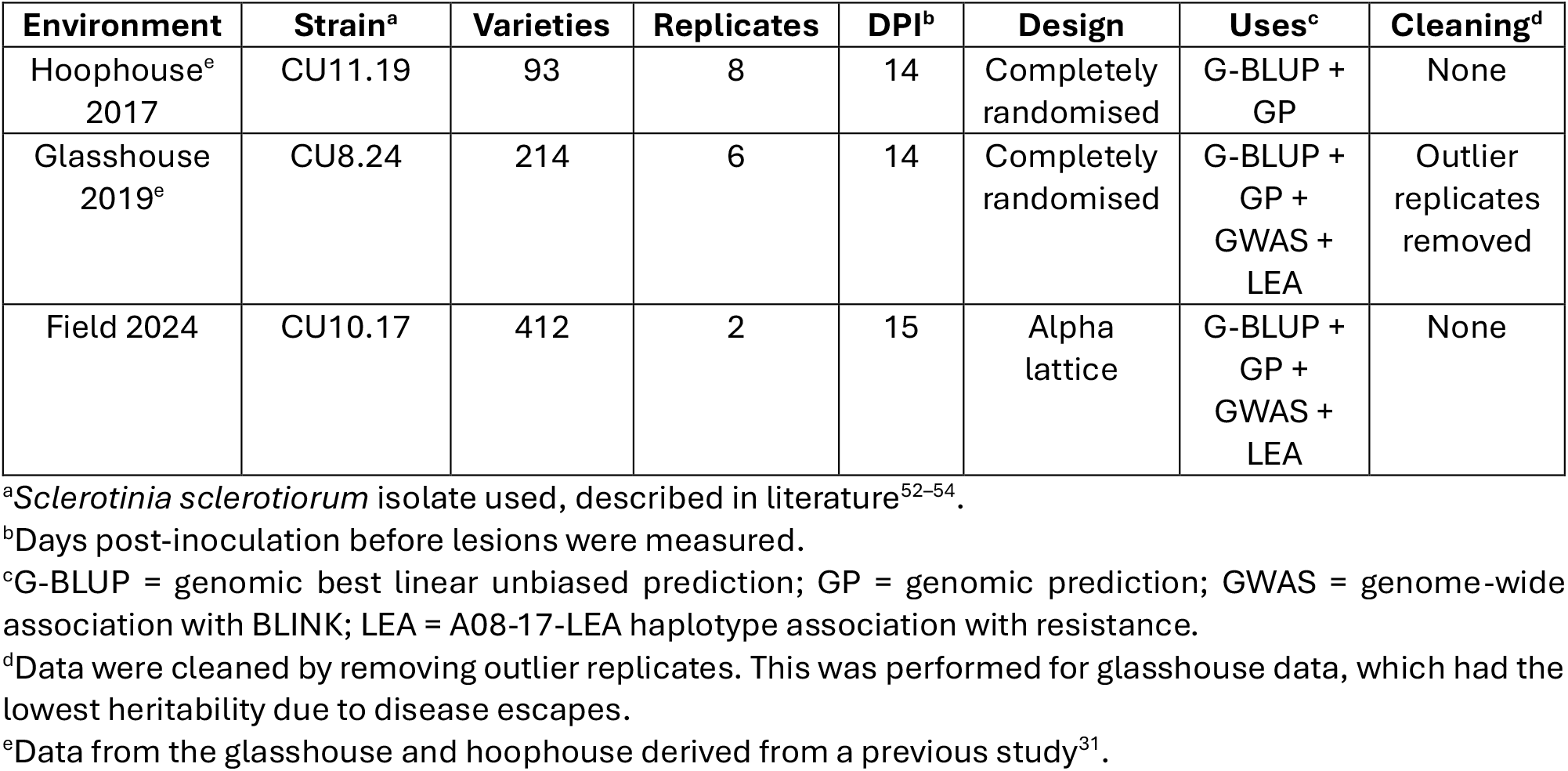
Overview of screening data sets.

| Environment | Strain<sup>a</sup> | Varieties | Replicates | DPI<sup>b</sup> | Design | Uses<sup>c</sup> | Cleaning<sup>d</sup> |
| --- | --- | --- | --- | --- | --- | --- | --- |
| Hoophouse<sup>e</sup><br>2017 | CU11.19 | 93 | 8 | 14 | Completely<br>randomised | G-BLUP +<br>GP | None |
| Glasshouse<br>2019<sup>e</sup> | CU8.24 | 214 | 6 | 14 | Completely<br>randomised | G-BLUP +<br>GP +<br>GWAS +<br>LEA | Outlier<br>replicates<br>removed |
| Field 2024 | CU10.17 | 412 | 2 | 15 | Alpha<br>lattice | G-BLUP +<br>GP +<br>GWAS +<br>LEA | None |

<sup>a</sup>*Sclerotinia sclerotiorum* isolate used, described in literature<sup>52-54</sup>.
<sup>b</sup>Days post-inoculation before lesions were measured.
<sup>c</sup>G-BLUP = genomic best linear unbiased prediction; GP = genomic prediction; GWAS = genome-wide association with BLINK; LEA = A08-17-LEA haplotype association with resistance.
<sup>d</sup>Data were cleaned by removing outlier replicates. This was performed for glasshouse data, which had the lowest heritability due to disease escapes.
<sup>e</sup>Data from the glasshouse and hoophouse derived from a previous study<sup>31</sup>.

Plants were grown and inoculated in the glasshouse and hoophouse as previously described^31^. Seedlings were grown in a glasshouse for the field experiment in UWA plant biology mix. At approximately the 3-4 leaf growth stage, seedlings were transplanted into a field plot. Before transplanting to the field plot, winter varieties were vernalised for 8 weeks at 4 °C in a cold room under growth lights set to an 8h:16h light:dark schedule. Plants were hand-watered and fertilised as required. They were inoculated individually at 30-50% flowering. Plants were inoculated using the method previously described^31^.

### Computational workflow management

Where indicated, computational work was executed using code hosted in GitHub repositories ccdm-bnpan (https://github.com/markcharder/ccdm-bnpan) and ccdm- bnpan-analysis (https://github.com/markcharder/ccdm-bnpan-analysis). Broad approaches are documented here with reference to these repositories for execution-level details and parameters.

### Genome assembly and annotation

The 22 new long reads genomes were assembled with Flye version 2.9.5 and Verkko version 2.2.1 (ccdm-bnpan/scripts/util/assemble_genomes_execute.sh). A single assembly was chosen for each genome based on N_50_ (Supplementary Table 1). Genomes were assembled from published Oxford Nanopore reads for Liho, Nugget and Tapidor (Supplementary Table 2) using Flye version 2.9.6 (ccdm-bnpan-analysis/scripts/liho- alleles-assembly.slurm).

Steps used in genome annotation are compiled in nextflow workflows in ccdm-bnpan. Before repeat annotation, genomes were scaffolded with RagTag 2.1.0 against the *B. napus* variety ZS11 reference genome^55^ (ccdm-bnpan/util/scaffold_genomes.sh). To avoid erroneous annotation of homoeologous genes as repeats, A and C genome scaffolds were annotated separately (ccdm-bnpan/workflows/assemprep.nf). Repeats were then annotated using panHiTE^56^ version 3.3.3 (ccdm-bnpan/workflows/panhite.nf).

Before gene annotation, >= 500 base pair contigs were softmasked with Repeat Masker^57^ version 1.4 using the library generated by panHiTE. These were concatenated with softmasked sub-genomes output by panHiTE (ccdm-bnpan/workflows/rnaseq.nf). *B. napus* RNA sequencing data optimised for genome coverage and mapping quality were then downloaded for each of the 22 genomes with Varus^58^ version 1.0.0 (ccdm- bnpan/workflows/rnaseq.nf). Braker3^59^ version 3.0.7.6 was used with these read mappings and proteins downloaded from OrthoDB version 11^60^ to annotate genes (ccdm- bnpan/workflows/braker3.nf). Translated coding sequences generated by braker3 were functionally annotated with InterProScan^61^ version 5.76-107.0 (ccdm- bnpan/workflows/interpro.nf).

### Graph construction and variant calling

A genome graph was constructed using the cactus 3.1.2^36^ sub-command ‘cactus- pangenome (ccdm-bnpan/doc/command-line/COMMANDS.md). This graph included un-scaffolded contigs for 22 new assemblies and 9 additional *B. napus* varieties^38,62–64^ (Supplementary Table 9). The *B. napus* variety X5A genome^38^ was used as a reference due to its exceptionally high quality. The ‘clipped’ graph was retained and the option ‘--hapl’ was used to generate haplotype information for haplotype aware mapping.

Illumina reads from 399 additional lines were mapped to the graph using haplotype sampling, with vg^65^ 1.70.0 (ccdm-bnpan/workflows/graphmap.nf) workflow. Briefly, kmc^66^ version 3.2.4 was used to generate a k-mer frequency file from read pairs. Reads were then mapped to the graph with haplotype sampling^67^ on-the-flye using vg giraffe^68^ (ccdm- bnpan/workflows/graphmap.nf). Alignments were filtered for quality with vg filter and variants were called from read mappings with vg snarls^69^ and vg call^70^ (ccdm- bnpan/workflows/graphcall.nf). Variants were called against the X5A reference genome^38^.

Resulting variant call format^71^ (VCF) files were processed with vcfbub version 0.1.2 to keep only top-level snarls (ccdm-bnpan/scripts/slurm/setonix_submit/vcfbub.run). The VCFs were then converted to binary call format (BCF) and merged in stages with bcftools^72^ version 1.23 (ccdm-bnpan/workflows/graphcall.nf).

Variants were then called directly from the GBZ^73^ formatted graph output by minigraph-cactus. After setting genotypes with conflicting traversals to missing (ccdm- bnpan/util/reannotate_vcf_conflicts.py), variants from the resulting VCF were merged with variants called from the graph with short reads using bcftools (ccdm- bnpan/doc/command-line/COMMANDS.md).

Variants called from the pan-genome graph and reads mapped to it were filtered using bcftools version 1.23-1-g638bee2d (ccdm-bnpan/doc/command-line/VARIANTS.md). Genotypes from that were heterozygous, had a GQ < 20 or DP < 2 were set to missing. Variants were then filtered to remove any with a minor allele frequency less than 0.05 and fraction of missing individuals of more than 0.1. Samples were then removed if they had more than 26% missing data. Across this subset, variants were then removed if they had a minor allele frequency < 0.05 and missingness rate >= 0.05. For all analyses relying on genotype data, biallelic SNPs from this quality filtered subset were used.

Further short and long read sequencing data were used to call variants in *B. napus* varieties Liho and Nugget. Short reads from accessions in Supplementary Table 4 were used to call variants in the same way as described above. Long reads were used to call structural variants against X5A (which carries “A08-17-LEA”) with sniffles^74^ version 2.8.0 based on minimap2^75^ version 2.28-r1209 mappings sorted with samtools^76^ version 1.11 (ccdm-bnpan-analysis/scripts/liho-alleles-svs.slurm).

### Estimation of genome-wide kinship and correlation with pedigree

Van Raden method 1^39^ was used to construct kinship matrices for population analyses with GCTA^77^ version 1.95 (ccdm-bnpan/doc/command-line/VARIANTS.md). Input genotype matrices were constructed for GCTA with plink^78^ version 2.0.0. Different levels of filtering for linkage disequilibrium were applied for different purposes using bcftools. For G-BLUP modelling, variants within 50 Kb of each other with R^2^ >= 0.99 were removed. For principal component analyses, variants within 100 Kb of each other with R^2^ >= 0.6 were removed.

A custom implementation of the Van Raden method was also used to estimate kinship between Australian cultivars, which was used for one of the selection scans and the heatmap in Figure 1 (ccdm-bnpan/doc/command-line/PEDIGREE.md). Population base allele frequencies were estimated based on the ecotypes of major founders of the Australian pedigree^14^ and multiplied by observed allele frequencies among sequenced founders. A kinship matrix was constructed for the Australian pedigree as per Van Raden method 1 with base R^79^ version 4.3.3 code, assuming these population base allele frequencies (ccdm-bnpan/envs/R/pedigree-env.R).

The kinship matrix constructed with GCTA was used to estimate correlation of genomic data with pedigree. A gamma matrix was first estimated by extracting kinship between global pedigree founders from the genomic data. Then, an additive genomic relationship matrix was constructed from the pedigree, conditioned on the known kinship among global founders (ccdm-bnpan/scripts/pedig-validate.r). Pearson’s correlation was then assessed for all pairwise kinships from the pedigree matrix and the genotype matrix, excluding the founders themselves. Correlations across the whole cohort and for pairwise comparisons between individuals and all other individuals in the pedigree were recorded.

### Selection scans

Three selection scans were implemented, based on a custom test for aberrant allele frequency shifts, Nei’s haplotype diversity^80^ and integrated haplotype score^81^ (iHS). The allele frequency test was conducted the following way. For cultivar *i* at SNP *k*, *g_i_* ∈ {0, 1, 2} defines the alternative allele count. Let *p* denote the alternative allele frequency at SNP *k* of the base population, which is a weighted sum of allele frequencies across assumed subtypes of the 22 founders contributing to sequenced cultivars. It is constructed using observed allele frequencies among sequenced founder subtypes, where *p_w_* = winter, *p_s_* = spring and *p_sw_* = semi-winter:

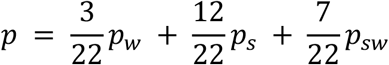

For cultivar *i*, the mean-centred genotype is *c_i_* = *g_i_* − 2*p*. The idea is to measure how much cultivars share an allele with one another and compare this with how much we would expect given the allele’s estimated frequency in founders. We do this by summing the product *c_i_c_j_* over all ordered pairs of *n* cultivars where *i* ≠ *j* in the following way:

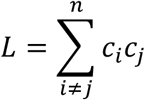

Then, we want to know what *L* would be at SNP *k* if it were a neutrally drawn allele from founders. The expected value of a cross-product at locus *k* under neutrality is:

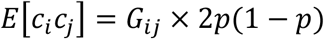

It follows that:

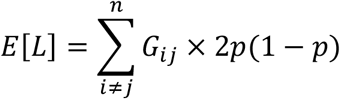

To derive the first term in the equation, we sum the off-diagonal elements of the kinship matrix constructed with Van Raden method 1, which define as *G_v_*. Therefore:

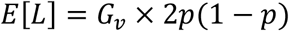

The AF test metrix, denoted simply as *T_AF_*, is then derived in the following way:

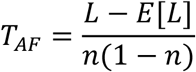

Bigger values of *T_AF_* denote bigger shifts in allele frequency from founders to descendants relative to what is expected given the estimated base population allele frequencies. To detect candidate selected regions, we identify the maximum value from 100 Kb sliding windows (overlapping by 50 Kb) and flag windows with a maximum value >= the 95^th^ percentile across all *T_AF_* scores for all SNPs.

The iHS score was calculated for 100 Kb windows using selscan^82^ version 3.3.0 across all Australian cultivars from inside and outside the documented pedigree. Nei’s haplotype diversity was also calculated across 100 Kb windows for sequenced Australian pedigree founders and pedigree cultivars, with the ratio of cultivar to founder haplotype diversity used as an assessment of selection. For iHS and Nei’s haplotype diversity, 100 Kb windows exceeding the 95^th^ percentile were again recorded as selection candidates (ccdm-bnpan/R/analysis/pedigree-sel.R).

### Genome synteny analysis

Genomic synteny dotplots (Figure 2 D) were constructed with custom code based on minimap2 alignments of un-scaffolded assemblies (ccdm-bnpan-analysis/scripts/liho- alleles-synteny.r). Structural variants were also called between scaffolded assemblies using syri version 1.7.1 and plotted with its utility plotsr (ccdm-bnpan- analysis/notebook/liho-alleles.md). To identify transposon insertions, a smaller graph was built from assembly scaffolds corresponding to X5A chromosome A08. Variants were called against A08-17-LEA carriers and founders of the Australian pedigree Haya, Westar and Zephyr. Using custom bash functions, genotypes from resulting VCFs were used to identify structural variants (>= 50 bp in at least on genotype) overlapping with transposon annotations from panhite (ccdm-bnpan-analysis/notebook/liho-alleles.md). Candidate genotypes were extracted with bcftools and aligned with mafft^83^ version 7.505. The regions surrounding candidates were also aligned with nucmer^84^ version 3.1 (option “— maxmatch”) and used to make synteny ribbon plots with gggenomes^85^ version 1.3.3 (ccdm-bnpan-analysis/scripts/liho-alleles-plotcands.r). To align the region in A08-17- LEA surrounding detected QTLs, we also used nucmer and made ribbon plots in the same way (ccdm-bnpan-analysis/scripts/sclero-resistance-lit-qtls.r). Complete gene absences from Mystic to Chikuzen and vice versa were defined as braker3 models completely within the bounds of insertions, deletions, highly divergent or not aligned regions called by syri (ccdm-bnpan-analysis/notebook/liho-alleles.md).

### Quantitative trait locus and linkage disequilibrium analysis

As a proxy for historical recombination, linkage disequilibrium (R^2^) was calculated between all pairs of SNPs within 100 Kb of one another using plink2 (ccdm-bnpan- analysis/notebook/liho-alleles.md). The results of this were used to evaluate average R^2^ across 100 Kb sliding windows in R (ccdm-bnpan-analysis/scripts/liho-alleles-plotld.r). This analysis was only performed on Japanese varieties from inside and outside the pedigree, since these varieties are not known to have been selected for low erucic acid and show normal levels of diversity across A08-17-LEA.

Genome-wide association analysis was conducted with BLINK^86^ version 0.1.0 on data from the glasshouse 2019 and field 2024 environments (ccdm- bnpan/R/analysis/genomodel-gwas.R). Genotypes used for this analysis were imputed for sites with missing data using beagle^87^ version 5.5. Raw phenotype data for both environments were normalised using a rank-based inverse normal transformation implemented in the R package bestNormalize version 1.9.2. Models were fit with and without population structure correction and quantile-quantile plots alongside inflation factors (*λ*) were used to choose an appropriate final model (Supplementary Figure 5) (ccdm-bnpan-analysis/scripts/sclero-resistance-qq.r).

For the glasshouse 2019 environment, phenotypes were means of transformed measurements. For the field 2024 environment, phenotypes were estimated marginal means from a linear model treating experimental block as a fixed effect. These were derived using the R package emmeans version 2.0.2. Each experimental block contained one replicate of each variety laid out in an alpha lattice design. Candidate QTLs were defined as SNPs meeting a false discovery rate^88^ threshold of 0.05 (scripts/sclero- resistance-gwasfilt.r). Consistency of effects across field and glasshouse environments was assessed with linear models including the SNP alleles and first three principal components from the PCA as covariates.

Additional QTLs were identified from the literature and projected onto the genomes of Mystic and Chikuzen. To do this, coordinates of *Sclerotinia* stem rot resistance QTLs on A08 in Darmor 4.1^89^ were taken from a recent QTL integration analysis^45^. The positions of Illumina 60K Infinium markers in scaffolded Chikuzen and Mystic genomes were then determined by mapping their sequences (downloaded from CropSNPdb^90^) with minmap2 and custom code (ccdm-bnpan-analysis/notebook/sclero-resistance.md). The approximate coordinates of QTL intervals from the literature were then defined as the closest Darmor 4.1 positions among Illumina SNPs. Positions of these SNPs based on minimap2 mappings were identified in the Chikuzen and Mystic genomes (ccdm-bnpan- analysis/scripts/sclero-resistance-lit-qtls.r).

### Genomic best linear unbiased predictor models and genomic prediction

G-BLUP^39^ models were used to assess heritability and genetic correlation across environments using the R package sommer^91^ version 4.4.5. Phenotypes were normalised as for the GWAS analysis but modelled across replicates rather than using expected marginal means. Models assumed the basic form:

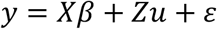

Where *β* contains fixed effects, including block for the field data, and intercept only for other environments. The genomic random effect 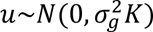 is conditioned on the kinship matrix described earlier and 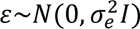, where *I* is an identity matrix. Heritability was estimated using single environment models with 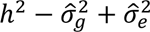. Genetic correlations across environments were estimated using a multi-kernel model with data from all three environments. The block fixed effect for the field environment was dropped and replaced with a fixed effect for environment and an unstructured covariance matrix was used to estimate the genomic effect across environments, defined as:

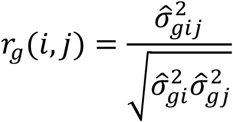

Genomic prediction was carried out for the field environment using the multikernel model fit to the same dataset with all field scores for Australian cultivars masked. Predictive ability was defined as Pearson’s correlation between predicted and observed scores (ccdm-bnpan-analysis/scripts/sclero-resistance-gp.r).

## Acknowledgements

This work was supported by resources provided by the Pawsey Supercomputing Research Centre’s Setonix Supercomputer (https://doi.org/10.48569/18sb-8s43) and Nimbus research cloud (https://doi.org/10.48569/v0j3-qd51), with funding from the Australian Government and the Government of Western Australia. MCD, YK, TN, NCW and LGK are funded by Centre for Crop and Disease Management, a co-investment between the Grains Research and Development Corporation (GRDC) of Australia and Curtin University (grant CUR00023). KKG is funded by Analytics for the Australian Grains Industry, a co-investment between the GRDC and Curtin University (grant CUR2210–005OPX). NCW is also supported by a research training programme (RTP) scholarship from the Australian Government.

## Author contributions

NCW contributed to study design, collection of data, analysis and manuscript writing and review.

KKG contributed to analysis and manuscript review.

TEN contributed to study design, collection of data and manuscript writing and review. YK contributed to collection of data and manuscript review.

LGK contributed to study design and manuscript review.

MCD contributed to research direction, establishment of the overarching conceptual framework, study design, development of most analyses and developed the first manuscript draft.

## Data availability

Data generated in this study will be made available in the national centre for biotechnology information sequence read archive upon publication.

## Notes

### Competing Interest Statement

The authors have declared no competing interest.

