## Supplementary figures for "Pedigree-reconciled pan-genomics reveals megabase-scale hitchhiking after a century of canola breeding"

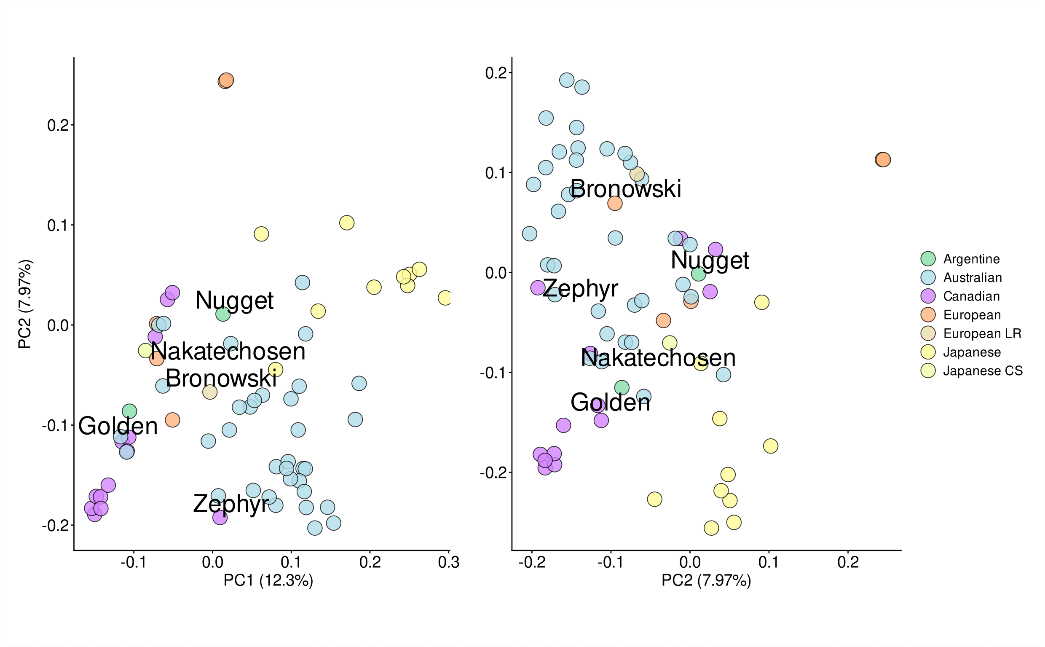

**Supplementary Figure 1. Principal component analysis based on genomic estimated kinship.** The plot to the right shows principal components 2 (7.97% variance) and 3 (4.8% variance).

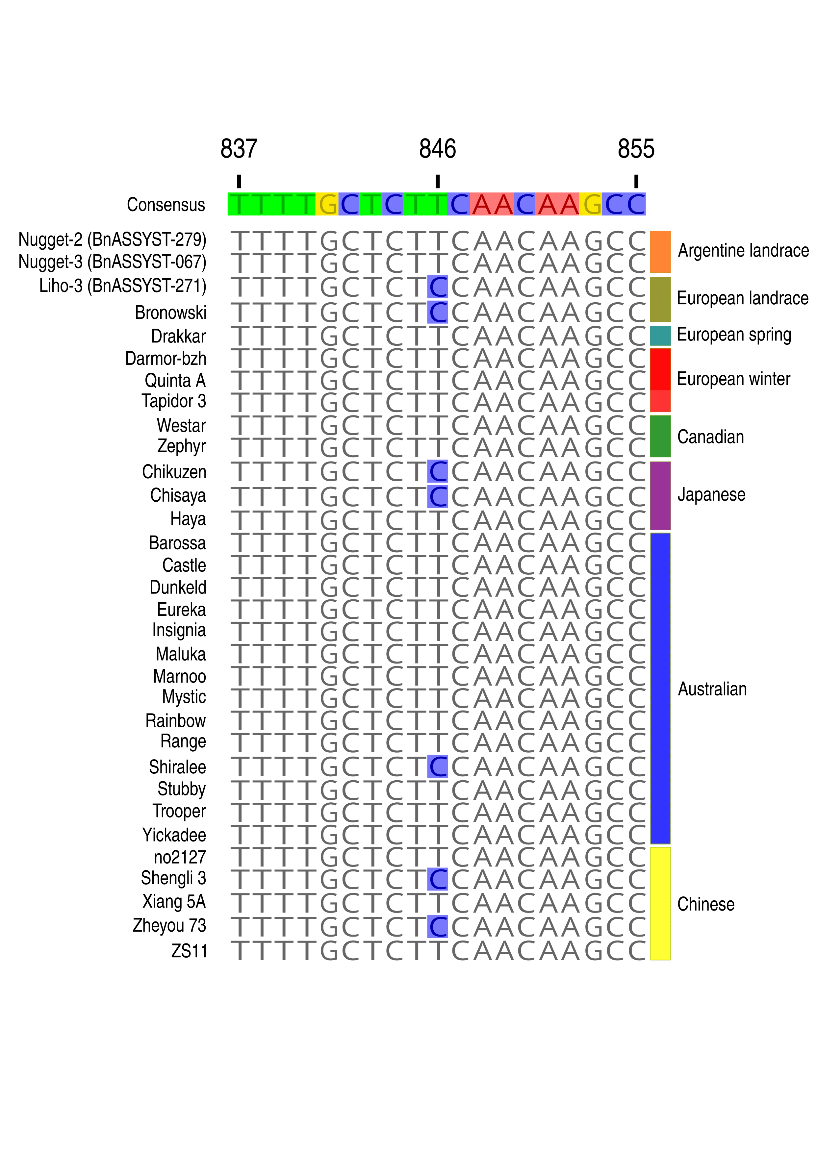

**Supplementary Figure 2. Alignment around the S282F substitution in *BnaA08.FAE1.*** The substitution is a C to T transition at position 846 in the gene. The high erucic acid allele is C, which is highlighted in blue as it differs from the consensus. *Brassica napus* type is to the right of the alignment.

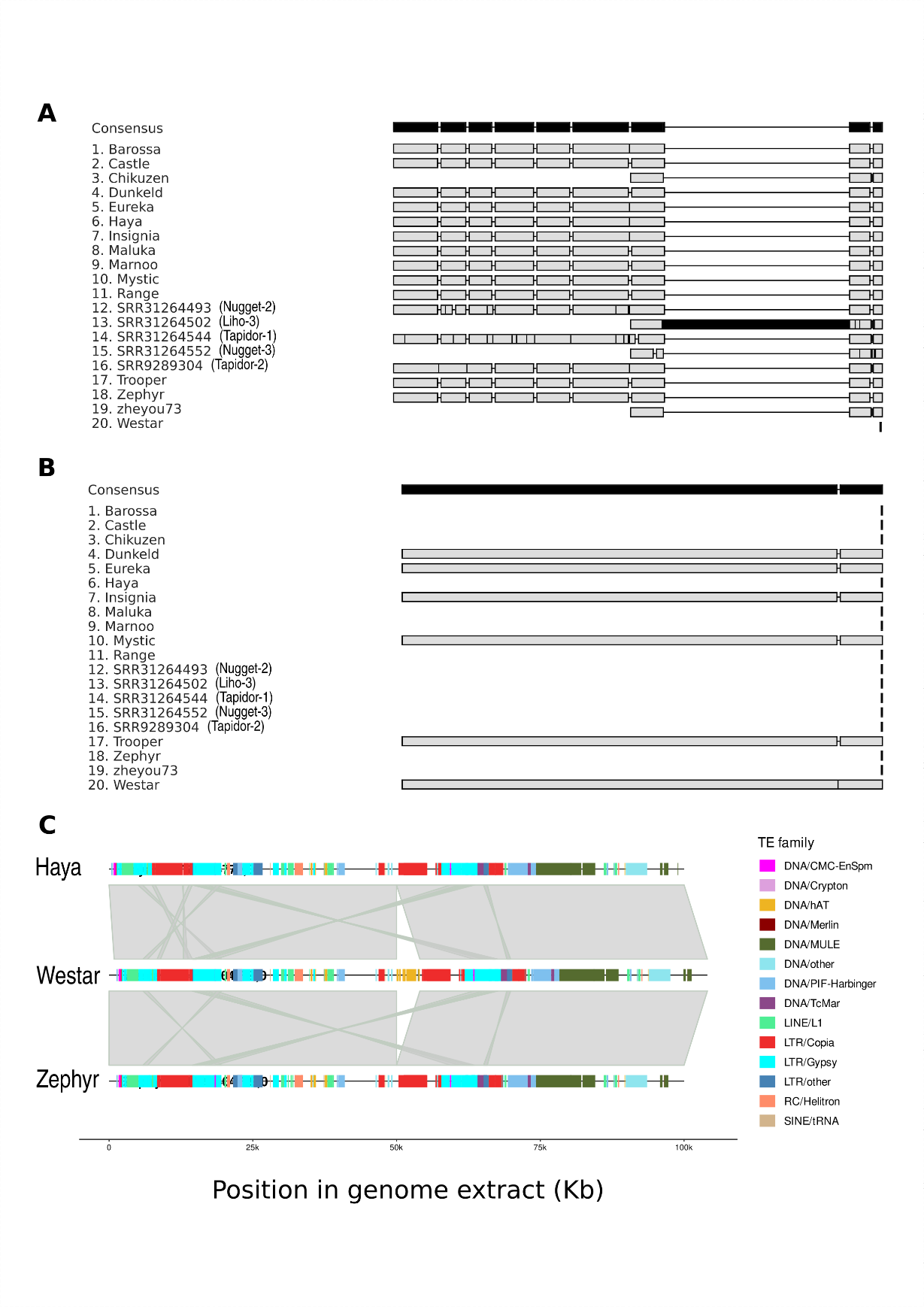

**Supplementary Figure 3. Alignment of alleles in A08-17-LEA in transposon insertion sites. A** Alignment of alleles across a LINE/L1 element insertion in Zephyr and Haya relative to Westar. **B** Alignment of alleles across a hAT transposon detected in Westar relative to Zephyr and Haya. **C** Showing the broader region for the alleles in **B**. A ~100 Kb region was extracted either side of the called structural variant and aligned with minmap2. The annotations represent repeat elements, including the hAT element inserted in Westar.

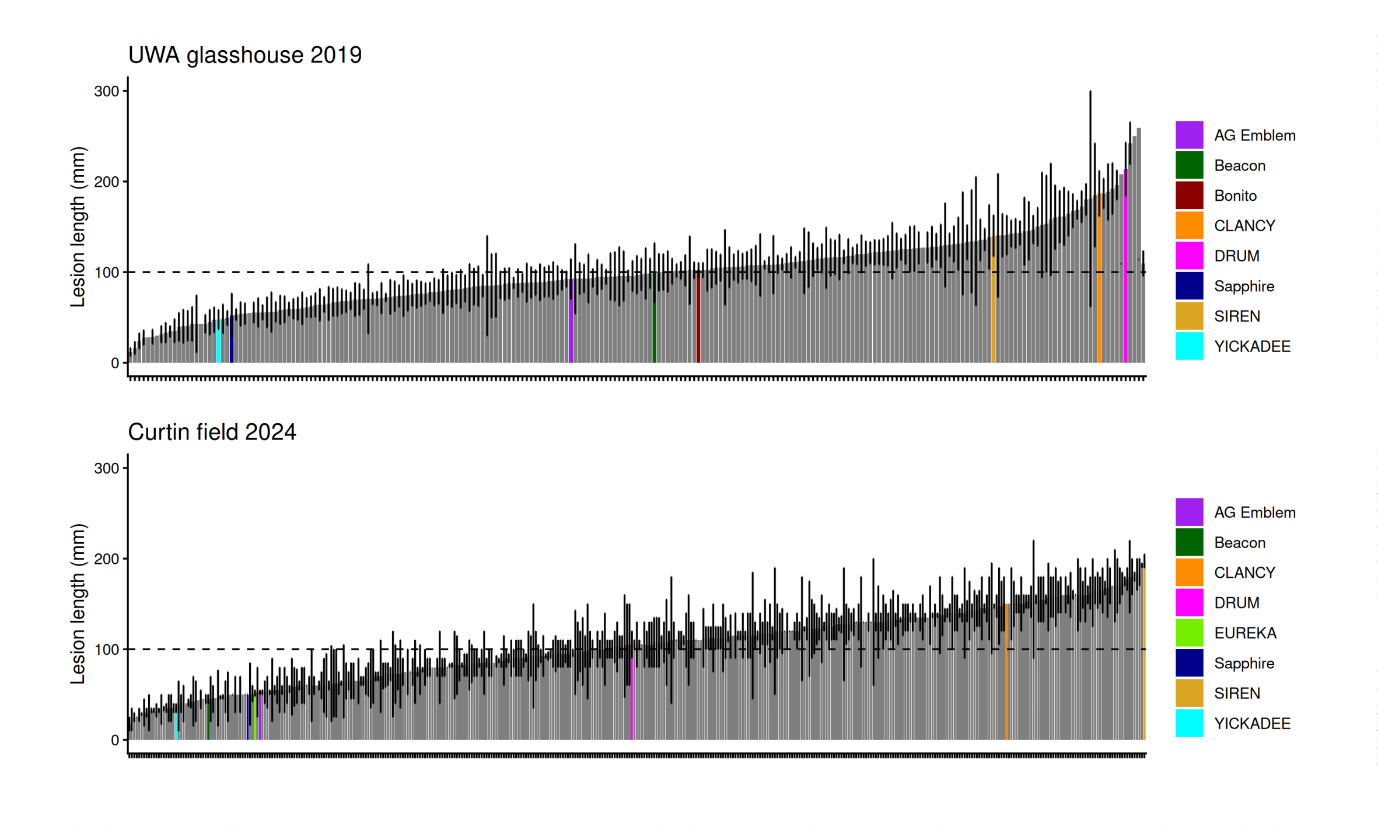

**Supplementary figure 4. Barplot showing lesion length (mm) in two environments.** Australian cultivars are marked out with coloured bars. The top panel is from a glasshouse screen with 6 replicates per variety and the bottom is a field plot screen with 2 replicates per variety. Yickadee and Beacon are descended from Chikuzen and Haya, which were found in this and previous studies to have similar levels of resistance.

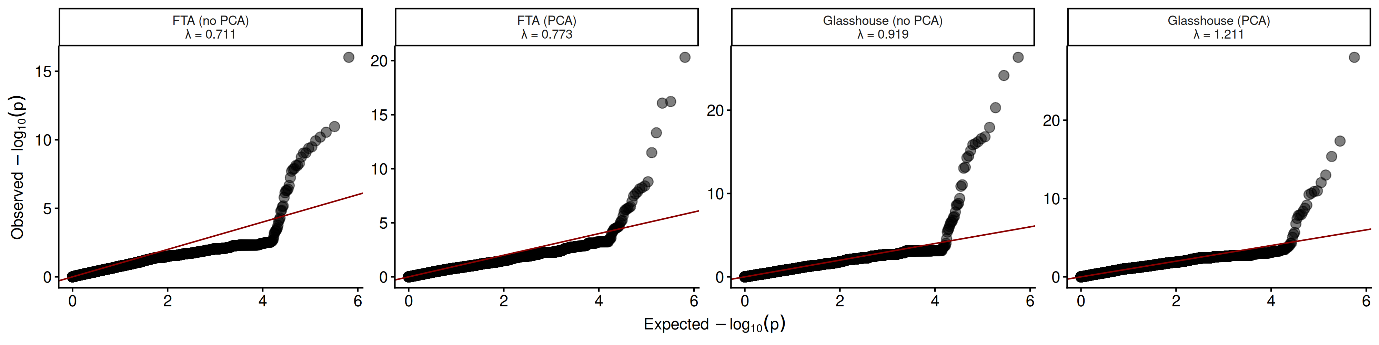

**Supplementary figure 5. Quantile quantile plots from genome-wide association testing.** The environments are FTA = field 2024 and Glasshouse = glasshouse 2019. The models chosen were field 2024 with principal component one (FTA (PCA)) fit as a covariate and glasshouse 2019 with no population structure correction (Glasshouse (no PCA)). Genomic inflation factors are given below. A value of below one indicates over-correction for population structure and a value above one indicates under-correction.
